# MassSpectrum Analyzer: An interactive platform for proteomic searching parameter refinement and peptide modification focused re-scoring

**DOI:** 10.64898/2026.06.22.733873

**Authors:** Kristian I. Karlic, Nichollas E. Scott

## Abstract

Peptide spectrum annotation is critical for the assignment of peptides and the localisation of modifications. While many existing tools provide spectrum annotation capacities, they often lack the flexibility required to allow bespoke spectral annotation of peptides containing multiple labile modifications or the accurate assignment of peptides in which fragmentation deviates from canonical patterns. In these cases, user-guided annotation is widely used to improve assignment completeness, however it typically does not integrate peptide scoring, making it challenging to assess the empirical improvement of the associated annotation and its impact on downstream false-discovery rate estimations. Here, we introduce an interactive annotation environment, the “MassSpectrum Analyzer”, which aims to streamline the exploration and analysis of modified peptides by enabling user–defined customisation with peptide scoring. Using (2–Aminoethyl)trimethylammonium carboxyl–derivatised peptides and glycopeptides as case studies we demonstrate the capacity of the MassSpectrum Analyzer to rapidly explore and allow the assessment of modified peptide datasets. By enabling direct assessment of the impact of user-guided choices on peptide scoring, we show how the detection of highly modified peptides can be improved through post-search integration of modification fragmentation information in a statistically robust manner. Similarly, by permitting comparisons of peptide ion intensities across spectra, we show that global fragmentation patterns can be quantified allowing the interrogation of trends that only become clear when spectra are assessed *en masse*. Combined, the MassSpectrum Analyzer streamlines the generation of publication-ready spectra and provides a means to assess how the inclusion of annotated features influences assignment scores.

## Introduction

The ability of protonated peptides to undergo predictable and informative fragmentation underpins bottom–up proteomics and the identification of peptides by tandem mass spectrometry (MS2) (1). Although the identification of peptides from MS2 events once required *de novo* interpretation by domain experts (2), database searching (3, 4) coupled to statistical filtering (5), using search tools is now the dominant means of peptide identification. By allowing the assignment of MS2 events as peptide spectral matches (PSMs), search tools underpin both qualitative and quantitative proteomics and are critically important for detecting and characterising protein modifications (6–10). As modified peptides can produce product ions in which the modification is intact, partially lost, or completely lost from the peptide backbone the MS2 events of these peptides can be substantially more complex than those of unmodified peptides (8, 11–13). This is exemplified by glycopeptides, which under collisional activation can yield peptide product ions with intact, partially lost, or fully lost glycans (14–20). Notably, chemical derivatisation strategies can also introduce chemical entities that alter fragmentation pathways in ways that deviate from those observed within unmodified peptides (11, 13, 21). By accurately accounting for modification-specific fragmentation patterns this can improve the confidence in PSM assignments and, in the context of database searching, enhance the identification of modified peptides (8). While improvements in computational approaches have enhanced the fields’ ability to identify modifications (7, 8, 10, 22, 23), understanding fragmentation characteristics through manual inspection remains an important quality-control approach within the field.

To date, several tools have been developed to facilitate spectral annotation (24–29), underscoring the importance of peptide fragmentation visualisation for the community. Beyond simply enabling the generation of publication–grade PSM annotations several provide additional specialised functionalities improving users ability to assess their data: For example the xiSPEC (26) and Annotater (24) tools support the annotation of cross–linked peptides; The Universal Spectrum Explorer tool (25) focuses on allowing the comparison of peptide fragmentation to predicted spectra or user provided spectra; while the recently developed Spectral Cruncher (29) facilitates *de novo* tag generation. While these tools allow visualisation of peptide fragmentation on an individual PSM basis, applying annotation information across datasets and quantifying fragmentation trends remains challenging, especially for modification specific fragmentation with teams typically extracting this information with custom scripts (30). As the field has long recognised, detailed analysis of MS2 events of modified peptides can reveal diagnostic ions and modification–specific fragmentation not observed in unmodified peptides (16, 31–34). For example, it was recently shown that the proximity of lysine acylation events to the N–terminus influences acylation immonium ion intensity, and that by optimising data acquisition to account for this, the confidence of acylation analysis could be improved (35). While we and others have shown incorporating characteristic fragmentation information can improve database searching (36, 37), an alternative approach is to use this information as an additional discriminator after database searching which can be achieved using peptide re–scoring.

At its core peptide re-scoring seeks to improve the segregation of potentially true identifications from false positives within proteomic datasets (38–41). Within re-scoring approaches, peptide identification is improved by the integration of information not utilised within the initial database search such as fragment ion intensity, ion mobility or retention time information (42). Incorporating this information can produce substantial gains in peptide identifications, in the case of immunopeptidomics, the use of Prosit-based re-scoring has been demonstrated to result in a up to a sevenfold increase in peptide identifications (43). Due to improving identification rates, re-scoring tools including Prosit/Oktoberfest (38, 44), MS2 Rescore (39), Inferys Rescoring (40) and MSBooster (41) have become integral components of modern proteomic pipelines. Although re-scoring approaches can substantially improve identification rates, they are generally best suited for unmodified peptides for which extensive training data has been generated (44). While recent efforts have attempted to address this limitation through the collection of training sets of modified peptides (45), not all modified peptides display similar fragmentation/LC or ion mobility behaviour. As a result, prediction accuracy for modified peptides is likely hampered due to their underrepresentation in training datasets and modification–specific fragmentation effects. This highlights the need for alternative re-scoring strategies that can be tailored to specific PTMs and fragmentation behaviours rather than relying on currently available intensity, mobility or retention time–based prediction models.

Here we present the MassSpectrum Analyzer, a spectral annotation platform designed to enable flexible and detailed interrogation of peptide fragmentation behaviour. The software supports annotation of standard fragment ion series while providing customisation through user–defined modification remainder ions, diagnostic ions, and glycan Y–ion handling within individual spectra. In addition to visualisation, the platform enables extraction of quantitative metrics, including annotated fragment ion counts and observed intensities, and allows assessment of how altering annotation parameters impacts scoring at both the single–spectrum and dataset scales. By enabling peptide re–scoring of datasets, we demonstrate how the identification of extensively modified peptides can be improved in a statistically robust manner. Together, these features make the MassSpectrum Analyzer a tool designed to facilitate mechanistic understanding of complex fragmentation behaviour while providing accessible large–scale data extraction and re-scoring capabilities without the need for custom scripting or computational expertise.

## Results and discussion

### Design of MassSpectrum Analyzer

Accurate interpretation of tandem mass spectrometry data is critical for confident peptide assignments in proteomics, particularly when analysing peptides that exhibit complex or non-canonical fragmentation behaviour. Herein, we outline the development of the MassSpectrum Analyzer, an open–source graphical interface for spectral annotation across proteomics experiments, with a focus on enabling user exploration and refinement of peptide annotations associated with complex fragmentation behaviour. While several annotation tools exist (24–29) allowing the assignment of standard ion series (a, b, c, c−1, x, y, z, z+1, d, v, w, Glycan Y-ions), the MassSpectrum Analyzer focuses on enabling the quantification of global fragmentation trends to allow user–guided refinement of annotations and rescoring. By allowing customisation of peptide annotations to include modification–specific losses, modification–associated fragment ions as well as satellite ions (w, v, d) and internal fragment ions (b_i_y_j_) this tool aim to streamline data interrogation of large datasets **(Figure 1)**. MassSpectrum Analyzer currently supports the analysis of data-dependent acquisition (DDA) experiments collected in centroid or profile mode from Thermo Scientific Mass spectrometers (.raw) or within the .mzML format (46) and while individual spectra can be manually assessed its primary function is to allow the visualisation of PSMs assigned from MSFragger (47), MaxQuant (48), MetaMorpheus (49), and Byonic (50), with broader compatibility enabled through the psm_utils Python package (51). Critically, annotation–associated information can be extracted across entire datasets, including annotation coverage, annotated total ion current (TIC %), matched ion counts, complementary ion pairs, counts of consecutive ion series, and ion intensities, as well as assessments of how the incorporation of user–defined features impacts re-scoring (as assessed by X! Tandem scoring (52)). Quantifying these metrics enabling systematic analysis of fragmentation patterns and trends **(Figure 1)**.

**Figure 1.**
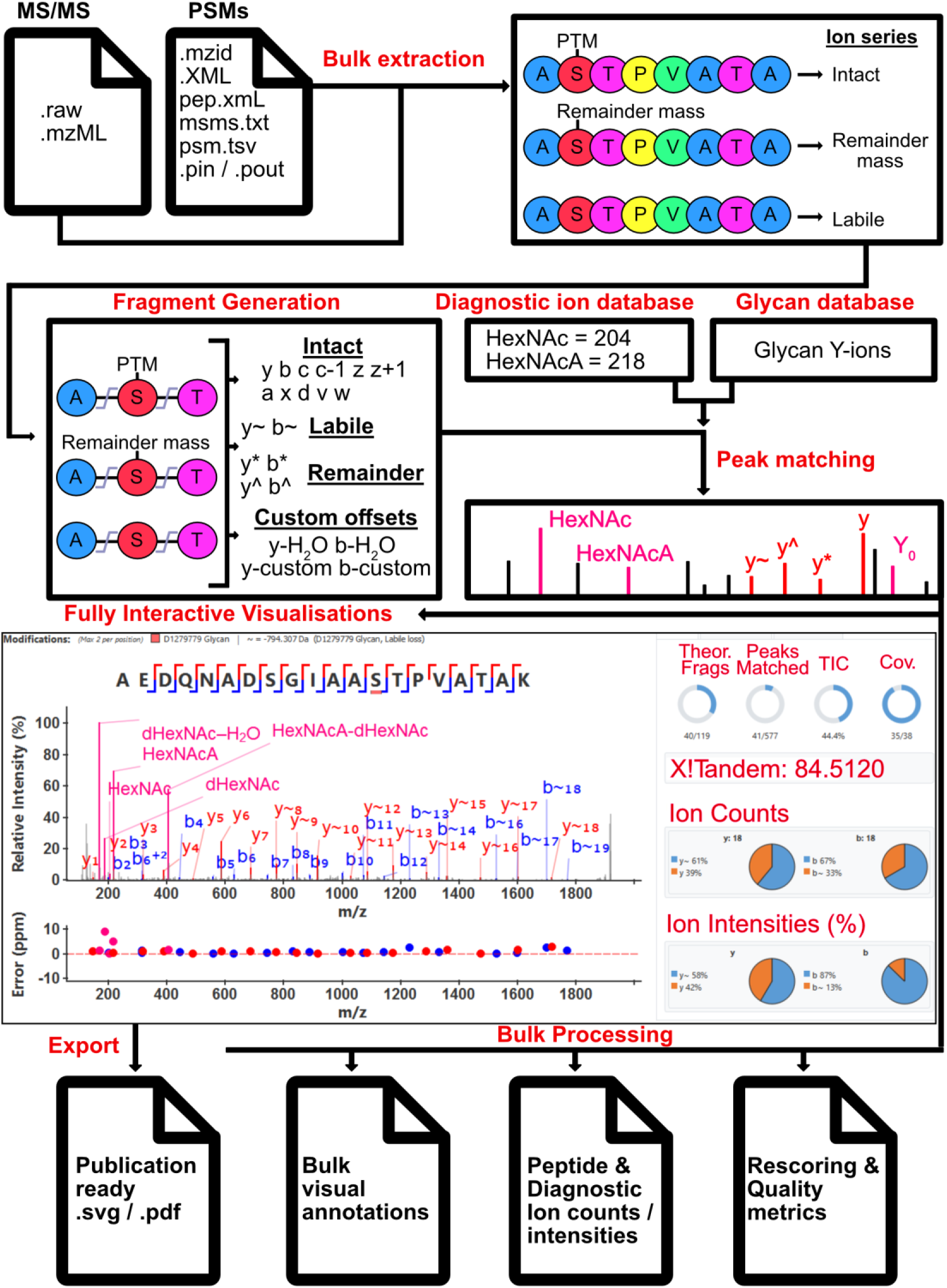
MassSpectrum Analyzer Flowchart. The MassSpectrum Analyzer processes .raw and .mzML datafiles, along with PSM identifications tables from commonly used search tools allowing the extraction and creation of saved experimental file optimized for PSM analysis. Users can define remainder masses, and modification lability in dedicated modification databases for re-use across experiments. The software matches theoretical peptide fragment ions, diagnostic ions, glycan Y-ions, against the PSM to produce an interactive spectrum. Users can export associated quality metrics including fragment ion counts/intensities as well as rescoring metrics from a single PSM or entire datasets *en masse*.

### User guided annotation allows improved peptide assignments

Confident peptide assignment requires accurate annotation of spectra, particularly for modified peptides, as modifications can produce distinct fragment ions that inform both peptide identification and the localisation of the modification (8, 11, 30, 53). These ions include fragments where the intact modification is retained on the peptide; fragments resulting from a partial loss of the modification (referred to herein as remainder fragments); and peptide fragments in which the modification is lost entirely, leaving no observable evidence of the prior modification. Within the MassSpectrum Analyzer *en masse* annotation of modifications from PSM search results can be undertaken allowing users to overlay custom fragmentation information on given modifications. Critically, this tool allows the visualisation of multiple fragmentation series associated with a given modification, rather than treating modifications as static additions a limitation of some currently available tools (24, 28), this can greatly improves assignment of observed features especially important chemoproteomics (11) studies. For example, we recently demonstrated that carboxyl–derivatisation of peptides with (2–aminoethyl)trimethylammonium (AETMA) results in the loss of 59 Da from AETMA under beam–type collision–induced dissociation (CID) (13). This AETMA-associated fragmentation has been previously reported (54) and results from the loss of trimethylamine leading to the formation of a three–membered aziridinium ring **(Figure 2A).** Using the MassSpectrum Analyzer, users can define additional annotation–associated information for a given modification applying modification rules across a dataset. The utility of this seen within the doubly AETMA–modified peptide ANSVKSALVNE[+84.1]YNVD[+84.1]ASR where by defining the loss of 59 Da from AETMA both modification events in this peptide are annotated, revealing that the majority of fragments correspond to ions with a single (−59 Da) or double (−118 Da) loss of trimethylamine (**Figure 2B–D**). By enabling users to define modification–associated features and apply them across a dataset this simplifies the annotation of large datasets.

**Figure 2.**
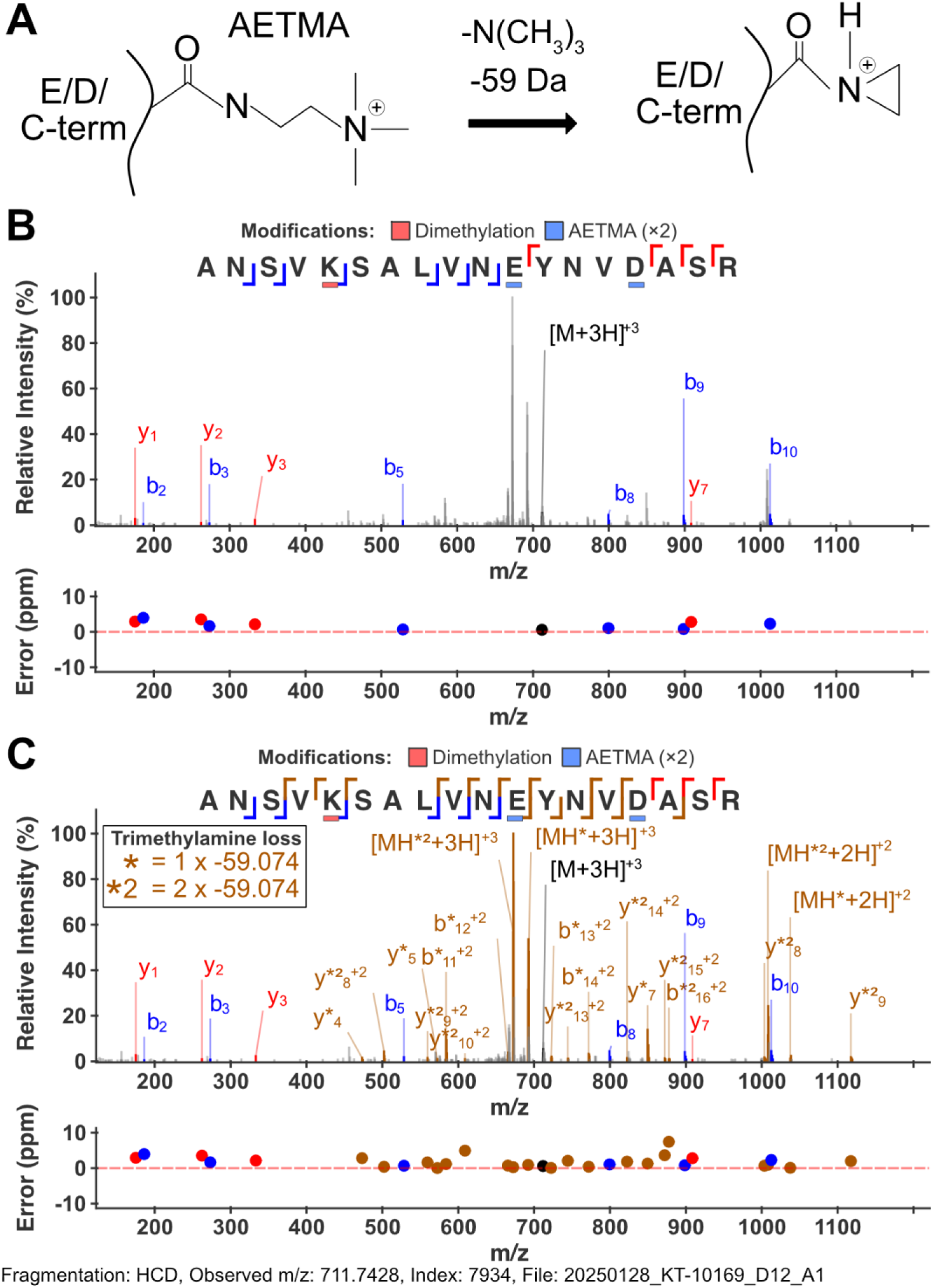
Peptide annotation in MassSpectrum Analyzer using combined non-labile and neutral loss fragmentation models. **(A)** Chemical schematic of AETMA fragmentation in response to collisional dissociation, whereby loss of trimethylamine occurs to form more stable aziridinium ring. **(B)** Spectral annotation with the MassSpectrum Analyzer of the *Acinetobacter baumannii* D1279779 peptide “ANSVK[+28.0313]SALVNE[+84.1054]YNVD[+84.1054]ASR” (NCBI Protein ID: AGH36670.1), containing E/D residues derivatised with AETMA and K dimethylation, using only b-, y-, and MH⁺ ion series. **(C)** Annotation with trimethylamine loss series * and *^2^ ions represent loss of -59.074 Da from AETMA modified fragments or x 2 -59.074 for double modified fragments, respectively.

An additional example where annotating multiple fragmentation series is advantageous is the assignment of glycopeptide PSMs in glycoproteomics studies. For N–glycopeptides, CID–based fragmentation is widely used due to its speed and optimal identification performance (30). As noted by several teams, the reducing N–acetylhexosamine (HexNAc) within N–linked glycopeptides is commonly retained on peptide ions as a +203.0794 Da modification to asparagine following glycan fragmentation (30, 55, 56). This property results in N–linked glycopeptide PSMs containing ions derived solely from glycan fragmentation (Y–ions) (57), peptide fragments retaining the reducing HexNAc, peptide fragments lacking any glycan–associated information, and glycan–associated oxonium ions (14). The MassSpectrum Analyzer supports the annotation of each of these ion series, allowing users to define ions associated with complete glycan loss (denoted y∼) as well as remainder masses associated with retention of the reducing HexNAc (denoted y^), as exemplified within the annotation of the glycopeptide N(+1216.423)VSWATGR (**Figure 3A**). Additionally, glycan Y-ions (57) are able to be annotated within MassSpectrum Analyzer using Symbol Nomenclature for Glycans (SNFG) (58), where users can define glycan compositions, and monosaccharide masses assigning them either a SNFG or simple letter notation can be used. The interface also enumerates annotation statistics within a given PSM, providing information on consecutive ion series (e.g., y_1_, y_2_, y_3_), complementary pairs (e.g., y₂/b₂), total annotated ion counts, and ion intensities to simplify the quantitative assessment of annotations (**Figure 3B**). Additionally, to improve the assessment of site localisation confidence three distinct measures of peptide sequence coverage are provided to users: I) Overall Coverage, defined as the total sequence coverage across all ion types (y, y∼, y^, y*); II) Intact Coverage, defined as the coverage from fragments retaining the intact modification (y); and III) Partial Coverage defined as the combined coverage from remainder and unmodified ions (y^, y*) (**Figure 3B**). By quantifying fragmentation coverage of a peptide in a modification–aware manner, these metrics aim to help researchers distinguish between confidence in peptide identification and confidence in modification localisation. Together, these features aim to provide users with additional metrics for assessing PSM and localisation quality beyond visualisation alone.

**Figure 3.**
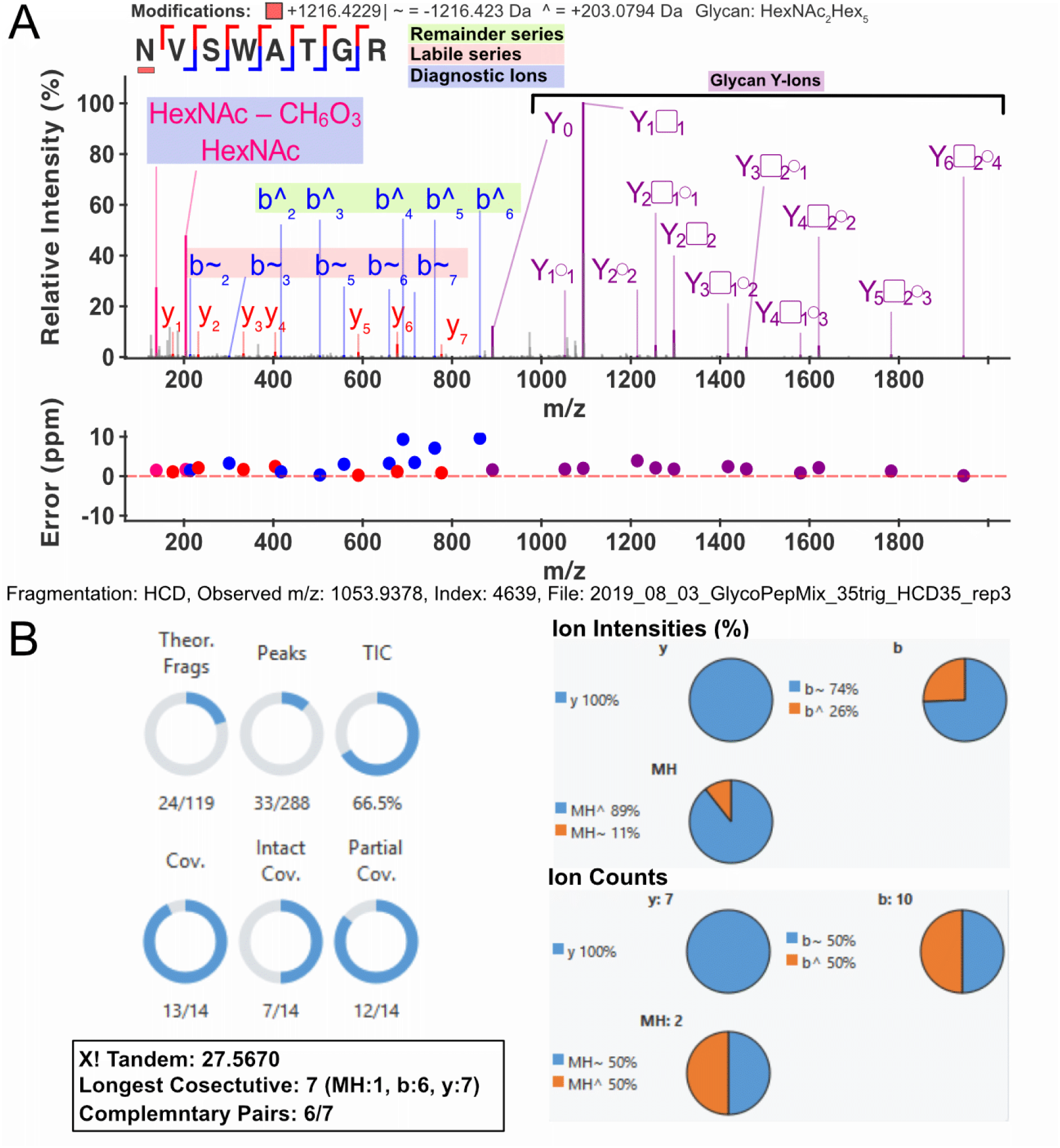
N-glycopeptide spectral annotation within the MassSpectrum Analzyer. Using data from Riley *et al*. (30) glycopeptide can be annotated and annotation metrics quantified. **(A)** Example human glycopeptide N(+1216.423)VSWATGR (Uniprot: P08571) annotated with y-/b-/Y-type ions, including remainder ions (+203.0794, denoted by ^), labile ions (+0, denoted by ∼), and oxonium ions. **(B)** Metrics calculated by MassSpectrum Analyzer for the annotated PSM. Metrics include: Theoretical Fragments annotated, Number of peaks annotated, Total Ion current (TIC%) annotated, Coverage (Overall peptide coverage – y/y∼/y^/y*), intact coverage (intact modification – y), and partial coverage (y/y^/y*), X! Tandem score, longest consecutive series of ions, complementary pairs, ion counts and ion intensities.

### Assessing Fragmentation Metrics Across datasets

Although detailed annotation of individual spectra is a powerful way to understand peptide fragment quantifying trends across large datasets is still a challenge for the field. Several existing tools allow the numeration of matched ions within PSM as well as the assessment of sequence coverage (24–29), however, expanding the set of metrics, including the *en masse* extraction of relative ion intensities of specific fragment ions, offers unique benefits for the analysis of multiple highly labile or unique modifications (8, 11–13). To address this the MassSpectrum Analyzer enables the extraction of several metrics to allow the quantification of fragmentation patterns including the relative abundance of fragment ions associated with intact, labile and remainder ions across datasets. To demonstrate the utility of tracking different fragment series for the analysis of modifications we assessed the impact of AETMA labelling density on observed loss of trimethylamine in response to CID (13). Using the MassSpectrum Analyzer bulk spectral extraction capabilities the relative intensity of y-/b-/a-/MH^+^ type ions across PSMs assigned to contain AETMA modified residues can be assessed at a dataset level (n=10,980). Quantification of trimethylamine loss associated with AETMA labelling density reveals that in peptides modified with two or more AETMA moieties, more than 50% of the observed y/b fragment ion intensity corresponds to ions associated with the loss of trimethylamine (**Figure 4A, Supplementary Table 3**). The high intensity of ions associated with trimethylamine loss also correlates with greater peptide sequence coverage when this loss is considered, compared with using intact–modification coverage alone (**Figure 4B**). Combined, these results demonstrate how global trends can be elucidated using the MassSpectrum Analyzer and shows that accounting for trimethylamine loss from AETMA improves the assignment of observed ion intensity and increases peptide sequence coverage across AETMA associated PSMs.

**Figure 4.**
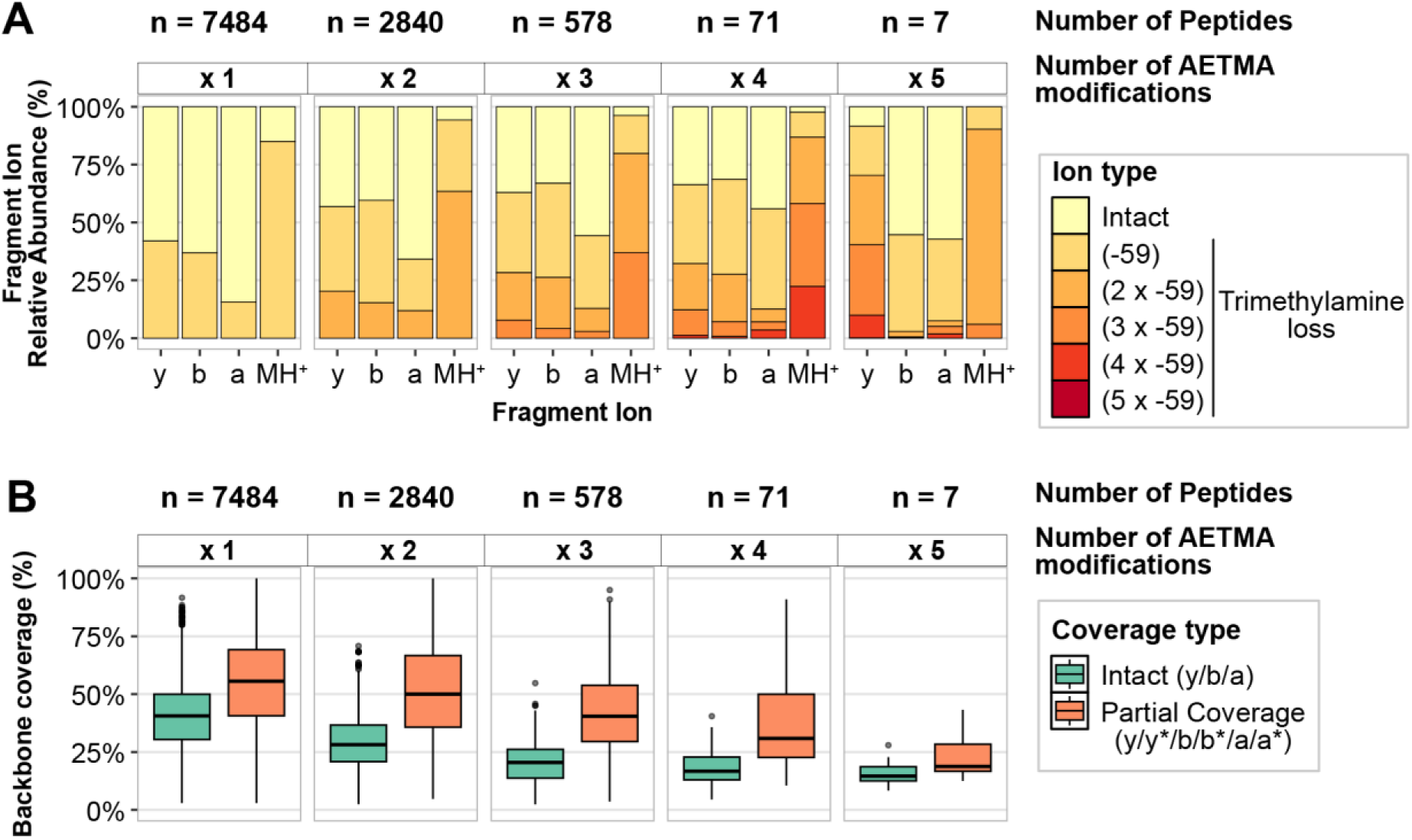
Trimethylamine loss associated with AETMA labelling density. Quantification of ions intensity and peptide coverage associated with AETMA modified PSMs identified using the MassSpectrum Analyzer. **(A)** The relative abundance of fragment ions for peptides labelled with increasing number of AETMA modifications, highlighting increasing proportion of fragment ion intensity of Trimethylamine loss **(B)** Comparison of backbone coverage of AETMA modified peptides associated with Intact (only considering intact modifications), and partial coverage (considering intact and Trimethylamine loss). All ions were matched within a 10-ppm tolerance.

To further highlight the capabilities of the MassSpectrum Analyzer in assessing trends across modified peptides, we analysed the published N-glycopeptide dataset of Riley *et al.*, which explored the impact of varying collisional energies on N-linked glycopeptide fragmentation (30). At low collisional energies (20% Normalised Collision Energy, NCE), fragment ions that retain the reducing sugar HexNAc (+203 Da) are more abundant than labile peptide ions (**Figure 5A, Supplementary Table 4**). As the NCE increases this pattern reverses as glycans are progressively stripped from the peptide during fragmentation consistent with previous reports identifying 35%–40% NCE as optimal for balancing peptide backbone fragmentation with N-glycan localisation (43). In addition to allowing the quantification of peptide associated fragment ions the MassSpectrum Analyzer also allows users to extract modification specific ions such as carbohydrate oxonium ions observed within glycopeptide PSMs. While this feature is also present in recently developed tools such as Glycounter (59), its inclusion here highlights the growing recognition that variation in modification specific fragment ions can provide additional supporting evidence of the stereochemistry of a modification such as glycan (60–62). Tracking the intensities of oxonium ions derived from N-glycopeptides containing HexNAc_4_Hex_5_NeuAc_2_ or HexNAc_4_Hex_5_ (**Figure 5B**) also demonstrates the unique HexNAc and NeuAc fragmentation profiles associated with different NCEs can rapidly be extracted and visualised. This functionality provides a framework for researchers to investigate fragmentation mechanisms and extract information modification specific ions that are currently challenging to achieve using available annotation tools.

**Figure 5.**
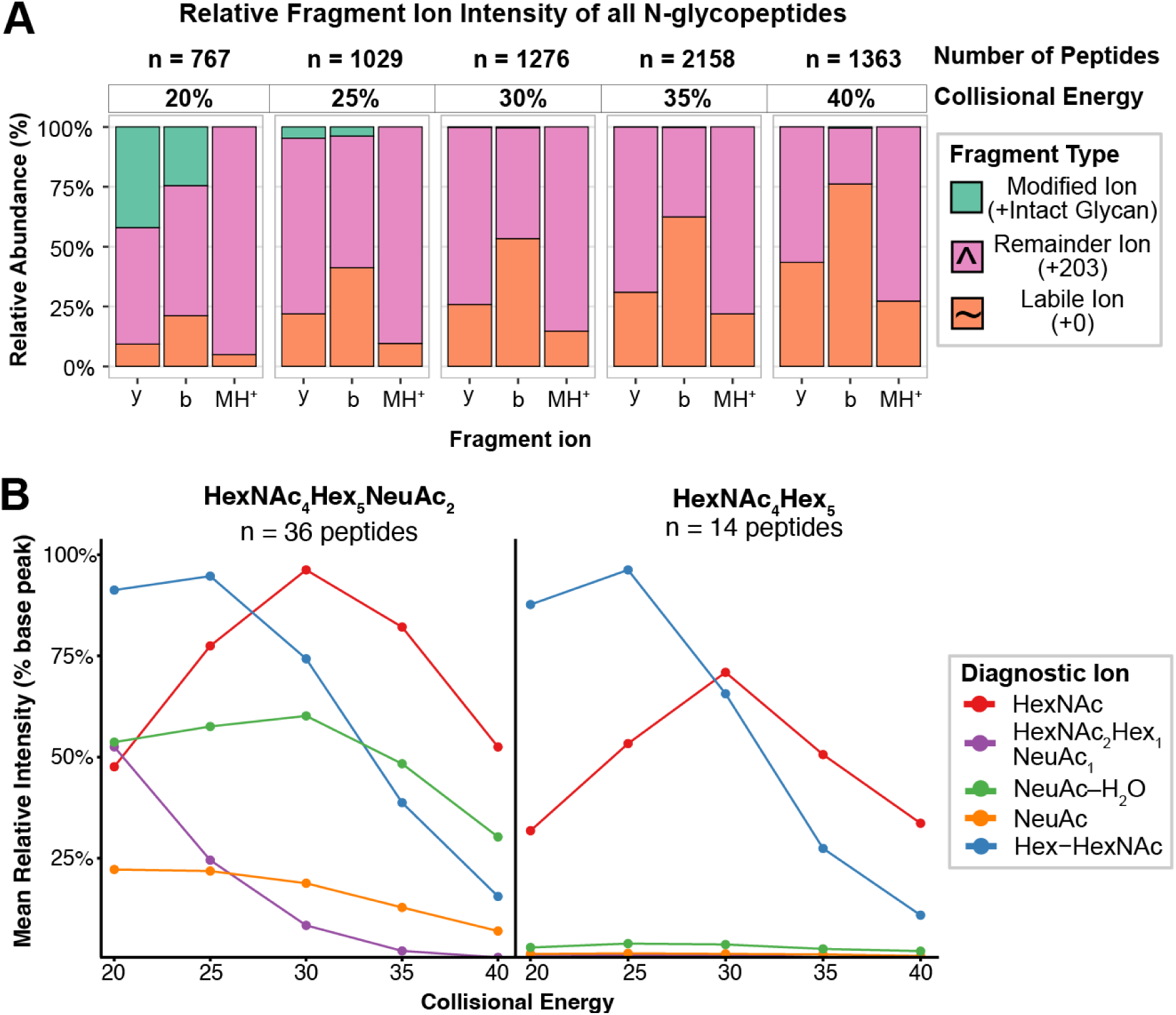
*En masse* analysis of N-glycopeptides trends. Glycopeptides PSMs matched ions, y-/b-/MH^+^ type ions, were extracted using the MassSpectrum Analyzer. **(A)** The relative ion intensity of different ion types including modified ions (+Intact glycan), remainder ions (+203), and labile ions (+0) from identified glycopeptides for y/b ions across increasing amounts of collisional dissociation. **(B)** Relative Intensity of oxonium ions for all glycopeptides modified with HexNAc_4_Hex_5_NeuAc_2_ or HexNAc_4_Hex_5_ across varying collisional energies. All ions were matched within a 10-ppm tolerance.

### Improving peptide assignments through user guided re-scoring

While the inclusion of user-guided information can improve the quality of PSM annotations, a key consideration currently unaddressed by available tools, is how to integrate additional assignment information into PSM scoring and downstream false-discovery rate (FDR) estimates within a given dataset. To address this the MassSpectrum Analyzer provides a suite of tools for rescoring of PSMs using X!Tandem-based scoring (52), as well as re-assessing a given datasets FDR using Mokapot, a Python implementation of the Percolator algorithm (63). By allowing users to tailor the fragment assignment information included within PSM rescoring the MassSpectrum Analyzer provides a means to assess impact of annotation in statically robust manner. To demonstrate the utility of rescoring we re-examined our AETMA modified peptides dataset (13) using MSFragger at 100% FDR in line with previous implementations of Prosit based rescoring (38, 64). Using an X!Tandem-based approach (52) incorporating b-and y-type ions alongside AETMA-specific loss of trimethylamine we assessed the number of identified PSMs at 1% and 5% FDR (**Figure 6A**, Supplementary Table 4). Rescoring resulted in an 11.8% and 4.2% increase in identified PSMs at the 1% and 5% FDR thresholds, respectively (**Figure 6B**), corresponding to an 8.6% and 3.2% increase in unique peptides identified at the same thresholds (**Figure 6C**). While rescoring produced higher absolute scores for both target and decoy PSMs, the separation between target and decoy distributions was improved relative to the original Hyperscore (**Figure S1A-C**). Consequently, several multiply modified peptides that failed to meet the 1% FDR threshold in the original analysis were successfully rescued (**Figure S1D, E**). This highlights the value of accounting for neutral losses when scoring chemically modified peptides. Ultimately, this provides users with a flexible framework for re-evaluating PSMs through the incorporation of expert, user-guided information.

**Figure 6.**
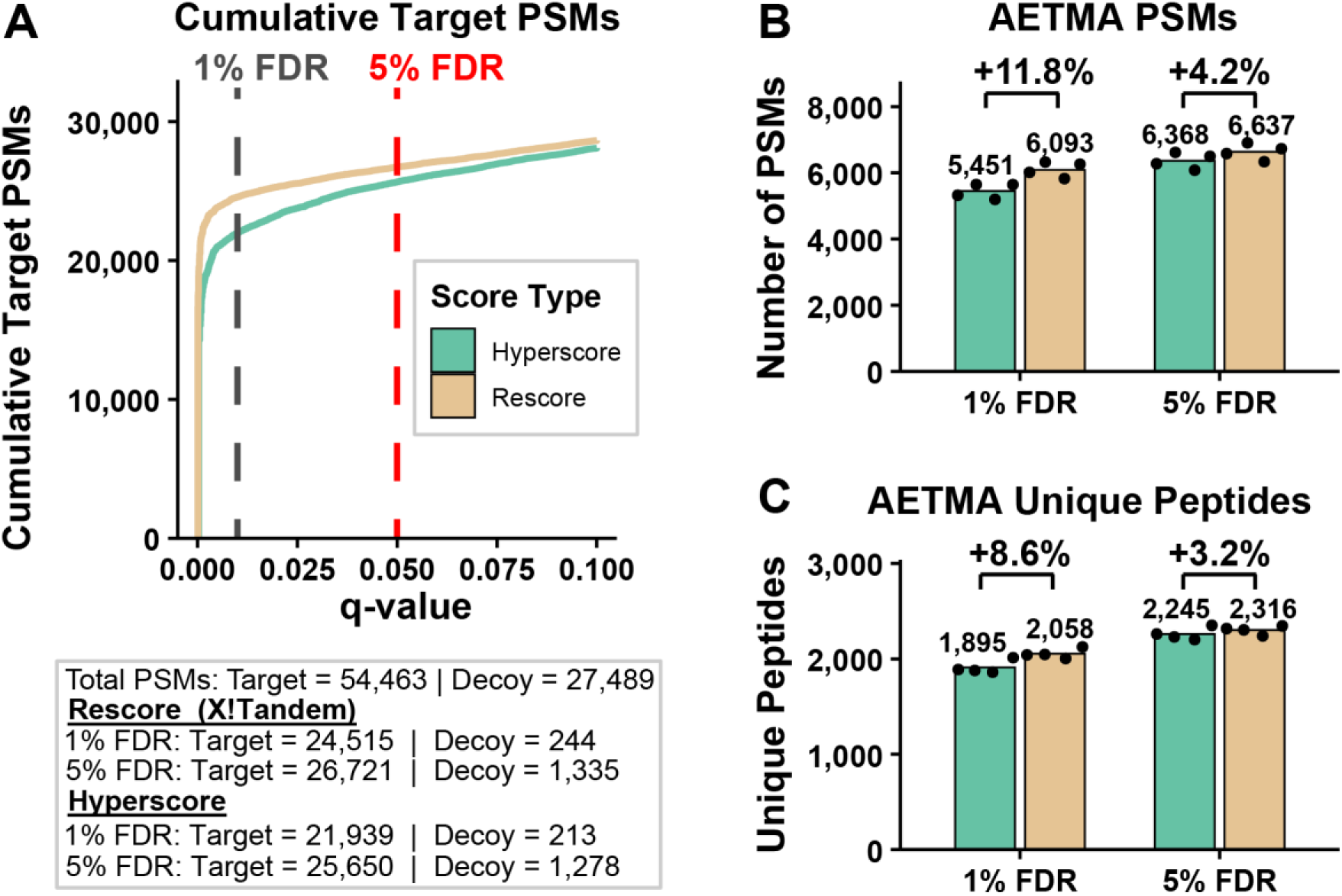
Rescoring. PSMs from *Acinetobacter baumannii* D1279779 peptides labelled with AETMA initially identified without FDR filtering from pepXML MSFragger outputs, were rescored and FDR analysis performed with Mokapot (63) within MassSpectrum Analyzer. For AETMA-labelled peptides a neutral loss of -59 Da for each modification (5 ppm fragment tolerance) was considered. **(A)** *A. baumannii* D1279779 PSMs were filtered at 1% and 5% FDR thresholds using both the MSFragger Hyperscore and MassSpectrum Analyzer Rescore (X!Tandem Method). **(B)** Number of target PSMs and **(C)** number of unique *A. baumannii* peptides identified after FDR filtering at 1% and 5% FDR.

### Improving peptide localisation confidence using peptide re-scoring

Confident localisation of post-translational modifications is an important aspect of peptide assignment, particularly for labile modifications such as O-glycans (14, 23, 30, 65). To improve user confidence in modification localisation, MassSpectrum Analyzer allows modifications to be repositioned interactively, with annotation metrics including X!Tandem score, TIC, complementary ion pairs and sequence coverage recalculated in real time. Using the *A. baumannii* BAL062 O-linked glycopeptide AKKEEAAQAGQDAAST(+843.31)AVADK from Tkalec *et al* (66) we observe the assignment of glycosylation to T15 (**Figure 7A**) yielding an X!Tandem score of 77.66. Repositioning the glycan to S14 (**Figure 7B**), in line with our previous work supporting Serine as the preferred site of glycosylation within *A. baumannii* (66), increases the X!Tandem score to 83.46, and results in longer consecutive ion series, higher TIC and greater sequence coverage. By allowing users to able to dynamically inspect how changes in modification localisation impact annotation metrics this feature aims to user help gauge site localisation assignments and the most likely site of localisation a given spectra. Finally, to streamline site of localisation evaluations the MassSpectrum Analyzer also allows rescoring of all residues to allow unbias localisation of potential modification (**Figure 7C**). These quantitative readouts enable users to make more informed, evidence-based decisions regarding modification localisation.

**Figure 7.**
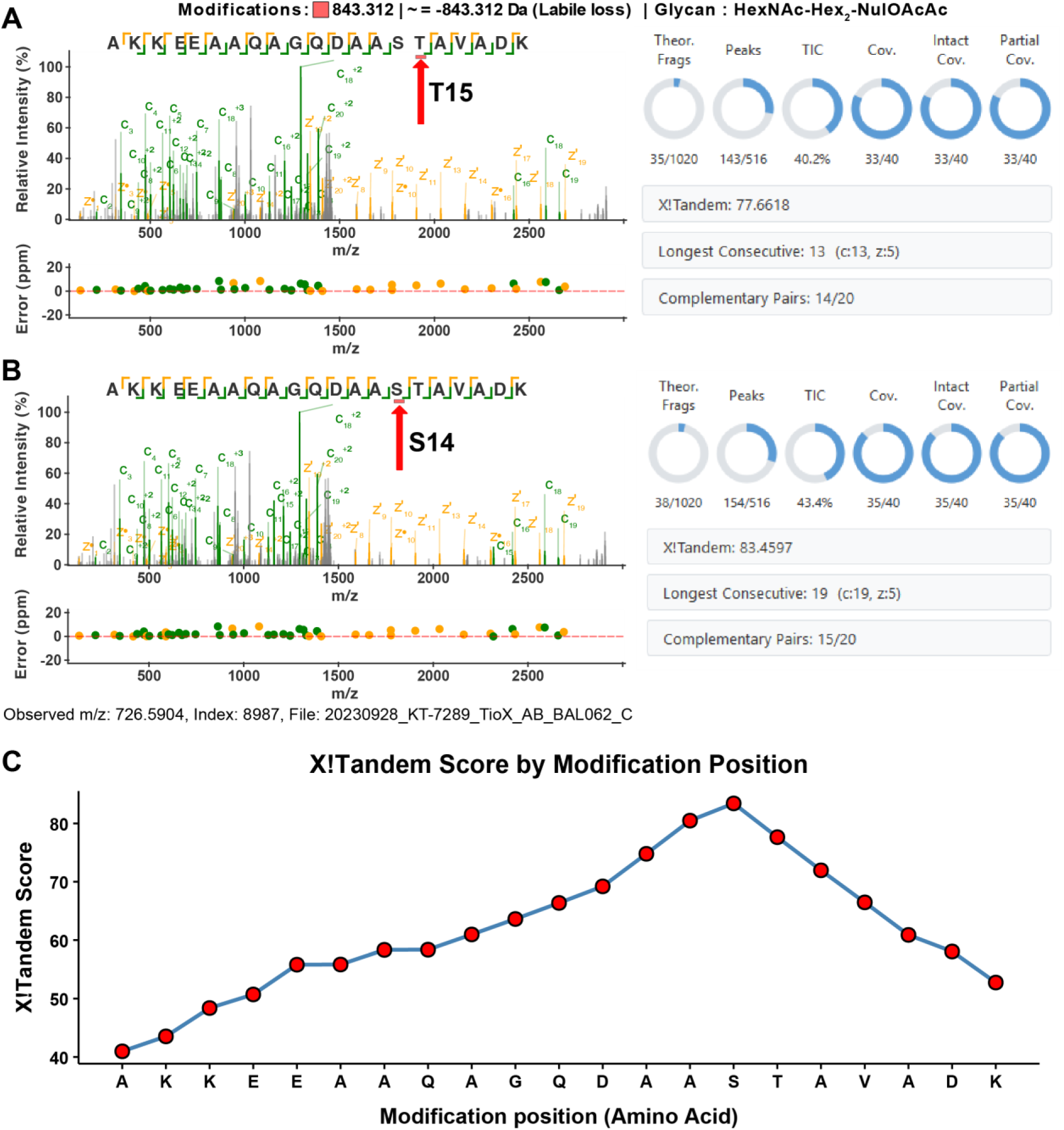
O-glycopeptide localisation example. The. MassSpectrum Analyzer allows for interactive repositioning of modification along the peptide triggering adaptive annotation and calculation of real-time annotation metrics to help users improve modification localisation. EThcD spectra and annotation information of an O-glycopeptide from *A. baumannii* BAL062, AKKEEAAQAGQDAASTAVADK (NCBI Protein ID: SBS22226.1) (66) with glycan **(A)** localised to T15, or **(B)** S14. **(C)** X!Tandem score for modification localisation of BAL062 glycan across all amino acid positions

## Discussion/Conclusion

Bottom–up mass spectrometry–based proteomics is an indispensable tool for the characterisation of modifications within peptides (67, 68). However, accurate annotation and enumeration of fragmentation patterns associated with modifications have largely been restricted to domain experts, owing to the challenges of analysing such data. Consequently, there is a critical need for tools that enable flexible visualisation and annotation of diverse datasets without requiring custom analysis pipelines, which can be inaccessible to non–expert users. The MassSpectrum Analyzer provides a flexible environment for peptide annotation, alongside tools to enumerate fragmentation information and extract metrics that are typically opaque at the dataset level (**Figure 1 and 3**). These capabilities allow researchers to better assess assignment confidence and to refine search results to more accurately reflect the interpretable spectral information. Using our recently published carboxyl–derivatized AETMA dataset (13) as well as an exemplar N-glycoproteomic dataset (30) we demonstrate the functionality of the MassSpectrum Analyzer and how this tool allows the exploration of modification at a PSM and dataset scale.

Our analysis of AETMA modified peptides demonstrates that for this chemical modification, the majority of peptide associated information corresponds to ions associated with the loss of trimethylamine (**Figure 4**), and integrating this loss into PSM scoring can improve the identification of AETMA modified peptides (**Figure 6**). This approach may prove valuable for other PTMs that produce characteristic remainder ions, where modification-specific fragmentation patterns are well-established but not always integrated into standard scoring (69–71). However, it is important to note that rescoring does not universally yield improvements, and the benefit appears contingent on the degree to which remainder ion contribute to the observed fragmentation. For example, within the chemoproteomic dataset of Yan *et al.* (11) which examines biotinylated cysteines peptides carrying a +463.237 Da modification that produces remainder ions of +152.095 Da and +180.101 Da, no improvement over the original search results was observed from peptide rescoring **(Figure S3A–E)**. Spectral examination of this dataset revealed that biotinylated cysteine remainder ions accounted for less than 10% of total ion intensity relative to the dominant y/b ion series (**Figure S2A-B**), supporting the conclusion that where remainder ions contribute only a limited proportion of observed ion intensity, rescoring is unlikely to provide meaningful benefit.

In addition, to enabling the assessment of chemically derivatised peptides, the MassSpectrum Analyzer provides a framework for the systematic interrogation of glycopeptide fragmentation, an area of growing importance within the proteomics field (59, 72–75). Our analysis of the N-glycopeptide dataset of Riley *et al.* (30) demonstrates that the tool can easily quantify relative contributions of intact, remainder and labile fragment ions within across collisional energies in the dataset (**Figure 5A**), and modification-specific oxonium ion profiles can be extracted *en masse* to reveal glycan-composition-dependent fragmentation trends (**Figure 5B**). Analysis of glycopeptides within the MassSpectrum Analyzer is also readily applicable to O-linked glycopeptides, where glycan lability under collisional energy produces predominantly labile ions and oxonium ions (14, 30). To illustrate this flexibility beyond eukaryotic N-linked glycosylation, we show that O-linked *A. baumannii* glycopeptides from strains D1279779 (76), and BAL062 (77) which utilise different glycan structures can annotated (**Figure S4**) (66), and localisation of O-linked glycans can be guided by the annotation metrics calculated during this process (**Figure 7**). Together, these examples demonstrate that the tool accommodates the diverse glycan compositions associated with bacterial glycosylation systems.

MassSpectrum Analyzer serves a distinct but complementary role to traditional proteomic search engines. Rather than performing PSM identification from raw data, it enables users to interrogate high-confidence PSM assignments to identify which annotation features drive, or fail to drive, confident scoring. Beyond this, the tool streamlines the extraction of diverse spectral quality metrics and quantification of matched ions, including fragment classes not routinely exported by existing search engines: intact, remainder, neutral loss, labile and diagnostic ions. This capability supports deeper exploration of fragmentation patterns at scale, as demonstrated by our analysis of previously published N-glycopeptide data acquired across multiple collisional energies (30). While MassSpectrum Analyzer already offers customisation not readily available in other tools, future development could extend this further; in particular, automated large-scale modification re-localisation across all possible modification states would be valuable enhancement, building on the manual adjustment functionality currently available through the interface (**Figure 7**).

Ultimately, by enabling users to extract a breadth of spectral information, MassSpectrum Analyzer facilitates a deeper understanding of complex fragmentation patterns and supports the refinement of PSM scoring in cases where current search tool fail to account for atypical fragmentation behaviour. We anticipate that the MassSpectrum Analyzer will serve as a valuable resource for the glycoproteomics and chemoproteomics communities, as well as the broader proteomics field, by facilitating deeper insights and improving the interpretation of complex mass spectrometry data.

## Methods

### Theoretical fragment ion generation

MassSpectrum Analyzer generates all theoretical fragment ion masses based on the ion types selected for fragmentation. This includes standard N-terminal ions (a, b, c) and C-terminal ions (x, y, z), satellite ions (w, d, v), and precursor (MH⁺) ion series across all positions along the peptide backbone within a specified charge range (+1 to a user-defined maximum charge). Peptide modifications are incorporated by applying their corresponding mass shifts to all fragments containing the modified residue. The user can also specify neutral losses (denoted *), remainder ions (denoted ^), or complete lability (denoted ∼) for modifications, enabling generation of these variant fragment ions. In line with the inhibition of electron fragmentation at proline (78) z-type ions are not calculated when the fragment sequence begins with proline, and c-type ions are excluded when the fragment ends with proline. To account of isotopic series up to four isotopes are considered and including in calculation for all generated fragment ions. Hydrogen transfer which is known lead to including c−1 and z+1 species is allowed and considered for ion calculations. Internal fragment ions (a, b) are generated from all subsequences longer than one residue and are calculated as singly charged species only.

Neutral loss species associated with amino acid composition including the losses of H₂O from residues S, T, E, D, NH₃ from R, K, Q, N and H₃PO₄ from phosphorylated S, T, Y or loss of SOCH₄ from oxidised M are enable by default yet the number of neutral losses can be defined by the user but is limited by the number of eligible residues within the fragment. In addition to the default ion series, the software supports user-defined fragment ion series with custom mass offsets, based on standard ion types (a, b, c, c−1, x, y, z, z+1, d, v, w, MH⁺). There is no explicit limit on the number of additional ion series that can be included, allowing for complex spectral annotation in cases with extensive fragmentation or neutral losses. These custom offsets can also be restricted to specific amino acid residues. All such ions are annotated alongside the selected base ion types, and any specified neutral losses can also be applied to these offset ions. Glycan Y-ions (57) can also be annotated for single spectra, where users can define glycan compositions within a glycan database (e.g. HexNAc(2)Hex(3)), and further define monosaccharide masses and assign generic SNFG symbol nomenclature (58, 79) for annotation using predefined Unicode shapes.

For each experimental peak, a mass window is defined using the user-specified ppm tolerance. All theoretical ions within this window are considered candidate matches. The fragment ion with the smallest absolute ppm error is selected as the primary annotation, while other candidates within the tolerance are retained as alternative assignments. Isotopic peaks are only matched if a corresponding monoisotopic peak has also been identified in the experimental data.

### Rescoring of PSMs

For each PSM the “Re-score” is calculated using the X!Tandem scoring method (52). The specific ion types (e.g. y/b/a) utilized in the calculation are user-specified, allowing for flexibility based on the fragmentation method employed. Furthermore, the maximum charge state considered for fragment ions is user-defined, with a default setting of +2. In addition to the primary rescoring metric, the software provides several complementary scoring features to assess match quality, including the percentage of annotated total ion current (%TIC), the identification of consecutive ion series, and the presence of complementary ion pairs. For *A. baumannii* D1279779 data derivatized with AETMA (PXD068858) and cysteine-modified data (PXD028853) (**Supplementary Table 1**), PSMs were initially identified using MSFragger (47). Precursor and fragment mass tolerances were both set to 20-ppm and fragment mass tolerance to 20 ppm. AETMA searches utilized a ArgC specificity (cleavage at R, not before P), while cysteine-modified data utilizing a tryptic specificity (cleavage at R/K, not before P), both with a peptide length range of 7–50 residues. For the AETMA dataset, variable modifications included methionine oxidation (max 2 per peptide) and AETMA derivatization (+84.10513 Da on glutamic acid, aspartic acid, and protein C-termini (max 5 per peptide). Carbamidomethylation of cysteine and dimethylation of lysine were set as fixed modifications. AETMA was further included as a mass offset in labile mode (8), with a fragment remainder mass of 25.03164 Da. For the cysteine biotinylation dataset (PXD028853), variable modifications included methionine oxidation (max 2 per peptide) and a +463.237 Da shift on cysteine (max 3 per peptide), specified as a labile modification with remainder ions of +152.095 Da and +180.101 Da. Fragment ion series included b-, and y-type ions. The resulting .pepXML output files, containing unfiltered target and decoy PSMs, were imported into MassSpectrum Analyzer. Rescored PSMs for both datasets were filtered at 1% and 5% FDR using mokapot (63) via the MassSpectrum Analyzer rescoring pipeline. For comparative consistency, the .pin files generated by MSFragger for each dataset were also processed through Mokapot for FDR filtering.

### Extraction of matched fragments

The re-scoring process within MassSpectrum Analyzer quantifies all user-selected fragment ions and diagnostic ions, which can then be exported in .csv long format for post-processing in environments such as R. To demonstrate this, N-glycopeptide HCD data at varying collisional energies were obtained from PXD017646 (30). Byonic viewer (50) was used to export PSM results as .csv files, which were subsequently imported into MassSpectrum Analyzer. Ion extraction was performed using the predefined “HCD-glyco” fragmentation method with the following parameters: standard ion types included MH, b-, and y-ions with remainder ions (+203.0794 Da), and labile ions (+0) variants. Oxonium ions extracted are described in **Supplementary Table 2**. All ions were matched within a 10-ppm tolerance. Visualisation of extracted ion fragmentation information was undertaken within R.

## Data availability

All R scripts used for analysis of outputs from MassSpectrum Analyzer is available at (https://github.com/Kristian-Karlic/MassSpectrum-Analyzer/tree/main/R_scripts_for_post_analysis). Data used for use cases of MassSpectrum Analyser were obtained from the following PRIDE accessions PXD068858 (13), PXD050066 (66), PXD017646 (30), PXD028853 (11), and specific files used for analysis is included in Supplementary data Table 1. MassSpectrum Analyzer is open source on GitHub (https://github.com/Kristian-Karlic/MassSpectrum-Analyzer), licensed under MIT.

## Supporting information

- Supplementary documentation containing; **(Table S1)** Files in Proteomics Identification Database (PRIDE) datasets, **(Table S2)** Oxonium ions matched during batch extraction , **(Figure S1)** Improved FDR filtering of AETMA labelled PSMs with MassSpectrum Analyzer, **(Figure S2)** Biotinylated cysteine peptide example, **(Figure S3)** Biotinylated cysteine rescoring shows no difference, **(Figure S4)** *A. baumannii* O*-*glycopeptide examples.
- **(Table S3)** Matched ion intensity of AETMA peptides (XLSX)
- **(Table S4)** Matched ion intensity of N-glycopeptides (XLSX)

## Acknowledgements

N.E.S and K.I.K are supported by an Australian Research Council (ARC) Future Fellowship (FT200100270), an ARC Discovery Project Grant (DP210100362) and a National Health and Medical Research Council Ideas grant (2018980).

## Author Contributions

K.I.K. – Data curation; Formal analysis; Investigation; Methodology; Validation; Visualization; Writing – original draft; Writing – review & editing

N.E.S. – Conceptualization; Funding acquisition; Methodology; Project administration; Resources; Supervision; Writing – review & editing

