## Supplementary_data for "MassSpectrum Analyzer: An interactive platform for proteomic searching parameter refinement and peptide modification focused re-scoring"

**Table of contents**

| **Supplementary Tables and Figures** | **Page** |
| --- | --- |
| **Table S1:** Files in Proteomics Identification Database (PRIDE) datasets | 3 |
| **Table S2:** Oxonium ions matched during batch extraction | 4 |
| **Figure S1:** Improved FDR filtering of AETMA labelled PSMs with MassSpectrum Analyzer | 5 |
| **Figure S2:** Biotinylated cysteine peptide example | 6 |
| **Figure S3:** Biotinylated cysteine rescoring shows no difference | 7 |
| **Figure S4:** *A. baumannii* O-glycopeptide examples | 8 |

**Additional Supplementary Tables**

**Supplementary Table 3:** Table_3_matched_ion_intensity_for_AETMA_peptides.xlsx

**Supplementary Table 4:** Table_4_Matched_ion_intensity_for_N_glycopeptides.xlsx

**Supplementary Table 1. Files in Proteomics Identification Database (PRIDE) datasets.**

| Figure | File Name | PRIDE |
| --- | --- | --- |
| 2, 4, 6, S1 | 20250128_KT-10169_D12_A1,  20250128_KT-10169_D12_A2,  20250128_KT-10169_D12_A3,  20250128_KT-10169_D12_A4 | PXD068858 |
| S2, S3 | 2021-07-20-KB-nofaims-SY65-1,  2021-07-20-KB-nofaims-SY65-2,  2021-07-20-KB-nofaims-SY65-1-labile.pepXML,  2021-07-20-KB-nofaims-SY65-2-labile.pepXML | PXD028853 |
| 3, 5 | 2019_08_03_GlycoPepMix_35trig_HCD40_rep3, 2019_08_03_GlycoPepMix_35trig_HCD35_rep3,  2019_08_03_GlycoPepMix_35trig_HCD30_rep3,  2019_08_03_GlycoPepMix_35trig_HCD25_rep3,  2019_08_03_GlycoPepMix_35trig_HCD20_rep3,  2019_08_03_GlycoPepMix_35trig_HCD40_rep2, 2019_08_03_GlycoPepMix_35trig_HCD35_rep2,  2019_08_03_GlycoPepMix_35trig_HCD30_rep2,  2019_08_03_GlycoPepMix_35trig_HCD25_rep2,  2019_08_03_GlycoPepMix_35trig_HCD20_rep2,  2019_08_03_GlycoPepMix_35trig_HCD40_rep1, 2019_08_03_GlycoPepMix_35trig_HCD35_rep1,  2019_08_03_GlycoPepMix_35trig_HCD30_rep1,  2019_08_03_GlycoPepMix_35trig_HCD25_rep1,  2019_08_03_GlycoPepMix_35trig_HCD20_rep1 | PXD017646 |
| 7, S4 | 20230928_KT-7289_TioX_AB_BAL062_C  20230314_KT-5941_D12_WC_B_Non_specific_glycan_trigger | PXD050066 |

**Supplementary Table 2. Oxonium ions matched during batch extraction.**

| Chemical Composition / Loss | m/z (Theoretical) |
| --- | --- |
| NeuAc−3H_2_O | 238.071 |
| NeuAc−2H_2_O | 256.0816 |
| NeuAc−H_2_O | 274.0921 |
| NeuAc (Intact) | 292.1027 |
| HexNAc−C2H_6_O_3_ | 126.055 |
| HexNAc−2H_2_O | 168.066 |
| HexNAc−CH_6_O_3_ | 138.055 |
| HexNAc−H_2_O | 186.076 |
| HexNAc−C2H_4_O_2_ | 144.066 |
| HexNAc (Intact) | 204.0866 |
| Hex−HexNAc | 366.14 |
| Hex | 163.0601 |
| HexNAc-Hex-NeuAc | 657.2349 |

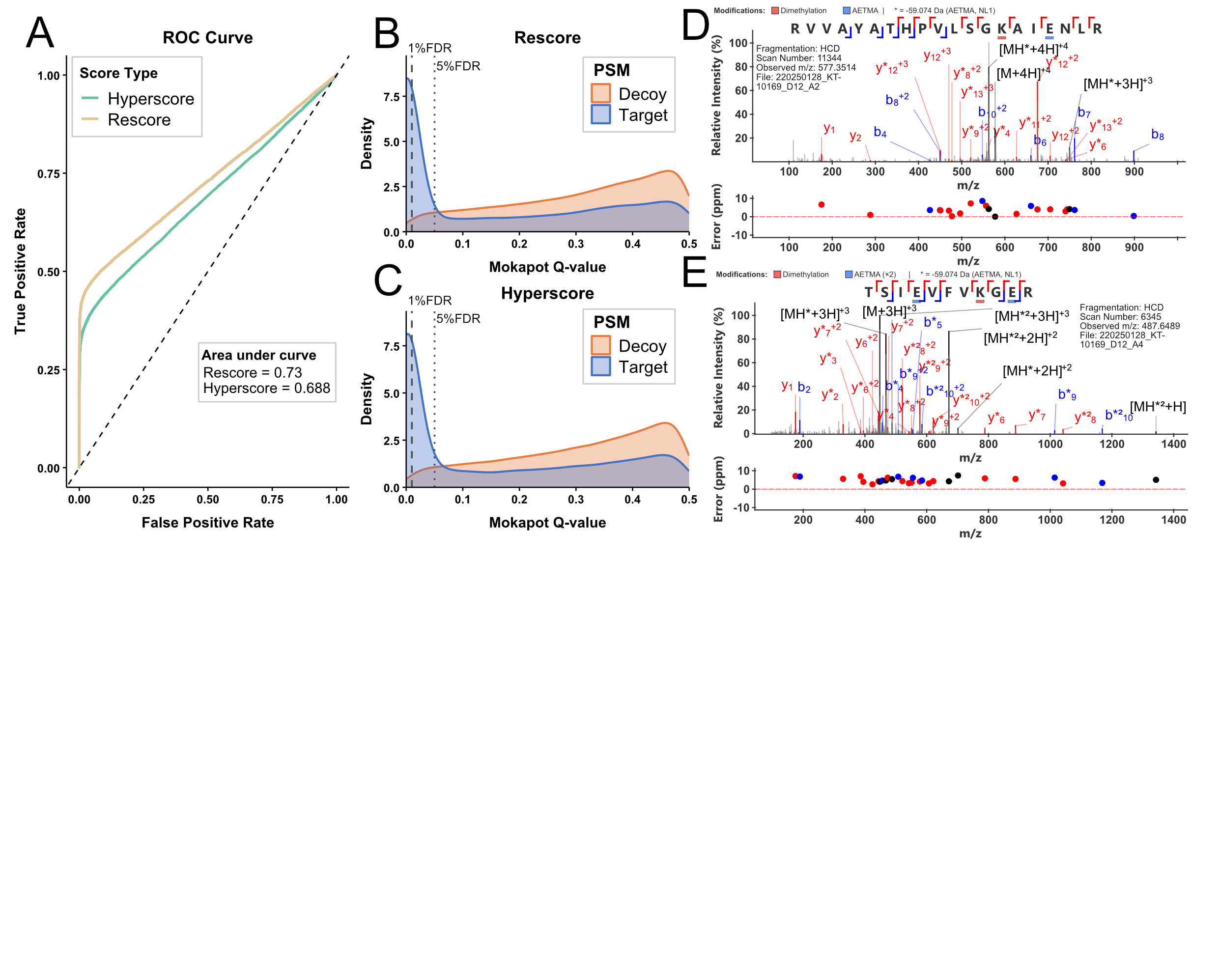

**Figure S1. Improved FDR filtering of AETMA labelled PSMs with MassSpectrum Analyzer. (A)** Receiver Operating Characteristic (ROC) curve showing that rescore improves target–decoy discrimination relative to hyperscore with a higher area under the curve. **(B-C)** Similar overall distribution of targets/decoys for rescore and hyperscore for Mokapot Q-value. **(D-E)** Unique AETMA labelled peptides RVVAYATHPVLSGKAIENLRR and TSIEVFVKGER unique to 1% FDR filtering of rescored PSMs.

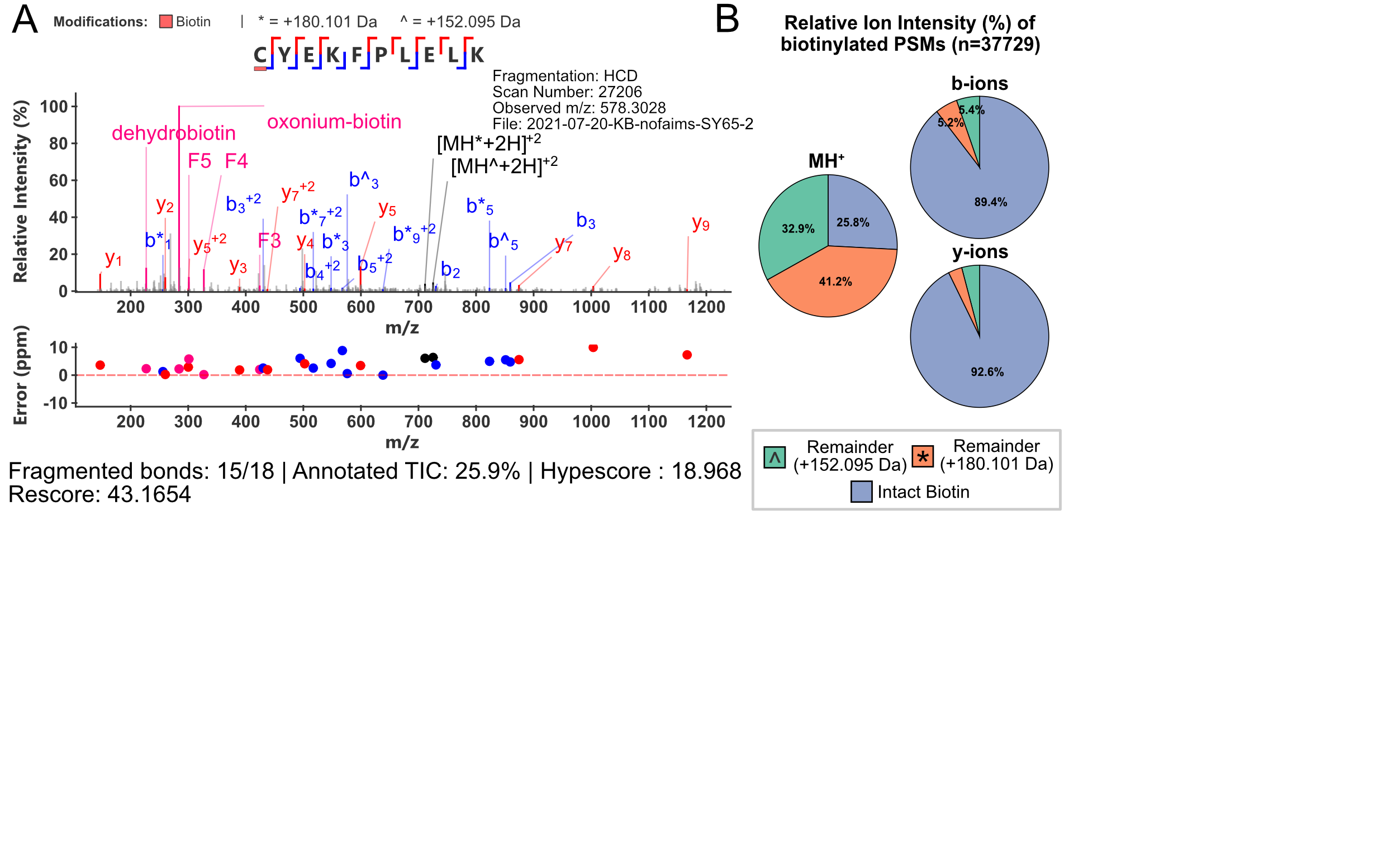

**Figure S2. Biotinylated cysteine peptide example. (A)** Spectral annotation for biotinylated peptide C[+463.2364]YEKFPLELK with annotation of remainder ion series 152.095 Da (^) and 180.101 Da (*), and diagnostic ions dehydrobiotin (227.085), oxonium-biotin (284.143) F3 (424.249), F4(327.185), F5 (301.169). **(B)** Relative Ion intensity of peptides fragments of biotinylated peptides for y/b/MH^+^ ions and remainder ion series (+152/+180). Fragment ions matched within 10-ppm fragment tolerance.

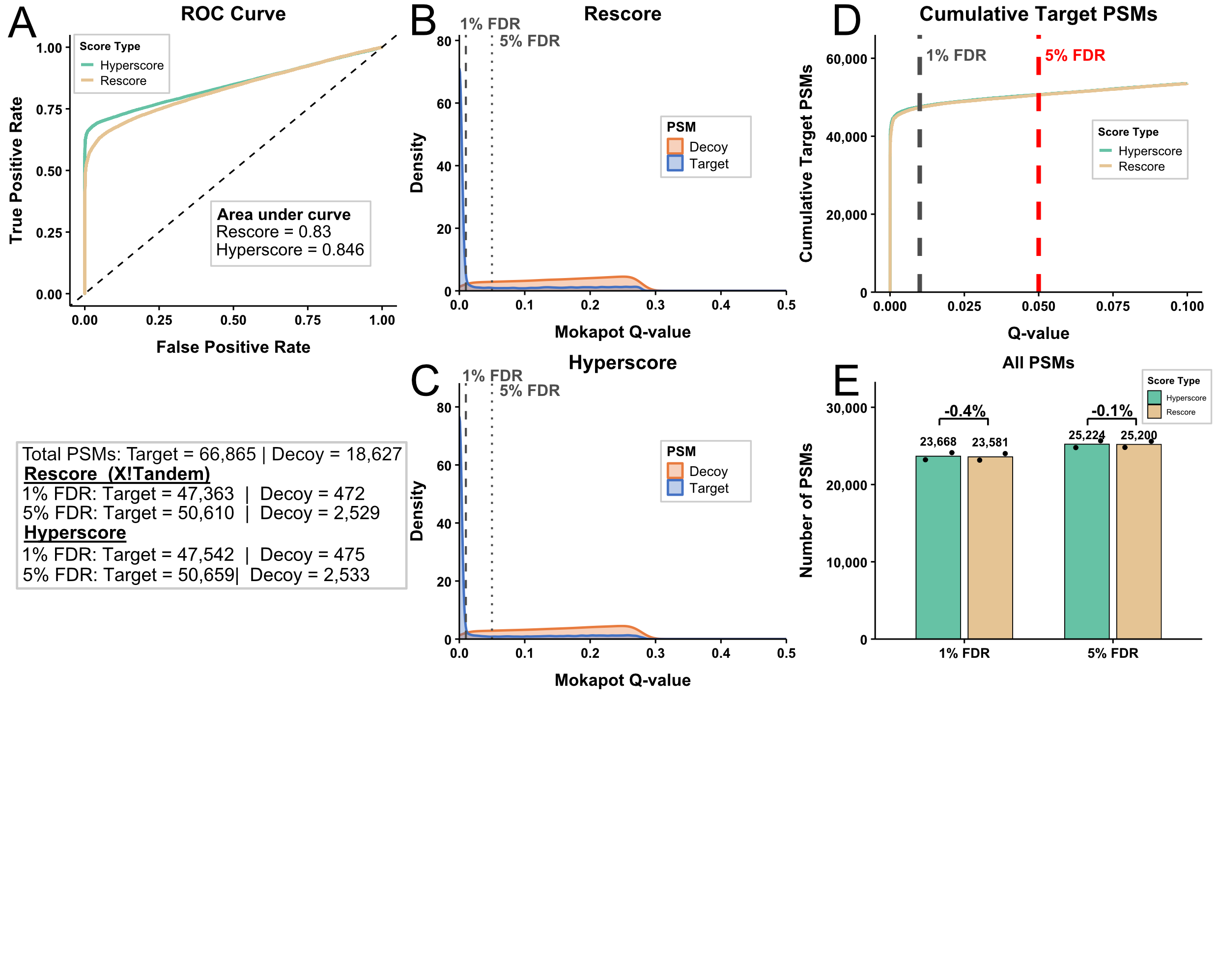

**Figure S3. Biotinylated cysteine rescoring shows no difference. (A)** Receiver Operating Characteristic (ROC) curve showing similar target–decoy discrimination relative to hyperscore. **(B-C)** Similar overall distribution of targets/decoys for rescore and hyperscore for Mokapot Q-value. **(D)** Similar target PSMs under 1%/5% filtering, and **(E)** negligible difference in mean number of PSMs between the two scoring methods for biotinylated PSM data. Fragment ions matched within 10-ppm fragment tolerance.

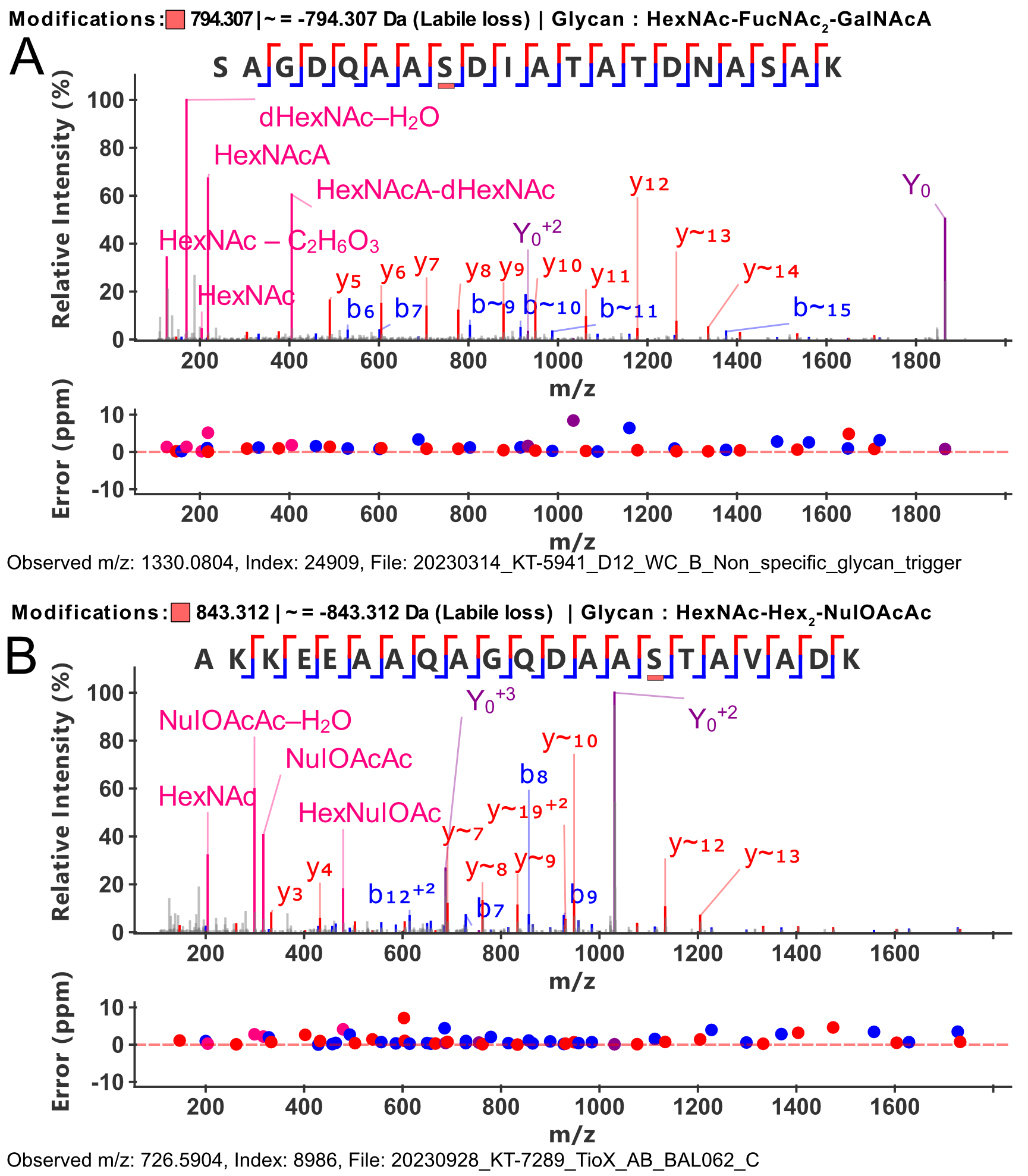

**Figure S4. *A. baumannii* O-glycopeptide examples. (A)** *A. baumannii* D1279779 O-glycopeptide modified with HexNAc-FucNAc_2_-GalNAcA (794.307 Da), **(B)** and *A. baumannii* BAL062 O-glycopeptide modified with HexNAc-Hex_2_-NulOAcAc (843.312 Da). b-/y-/Y- are matched with a 10-ppm tolerance.
